# Mavacamten stabilizes a folded-back sequestered super-relaxed state of β-cardiac myosin

**DOI:** 10.1101/266783

**Authors:** Robert L. Anderson, Darshan V. Trivedi, Saswata S. Sarkar, Marcus Henze, Weikang Ma, Henry Gong, Christopher S. Rogers, Fiona L. Wong, Makenna M. Morck, Jonathan G. Seidman, Kathleen M. Ruppel, Thomas C. Irving, Roger Cooke, Eric M. Green, James A. Spudich

**Author notes:** These authors contributed equally to this work (Robert L. Anderson, Darshan V. Trivedi, Saswata S. Sarkar). These authors contributed equally to this work (Eric M. Green, James A. Spudich). Correspondence (J.A.S); (E.M.G.).

## Abstract

Mutations in β-cardiac myosin, the predominant motor protein for human heart contraction, can alter power output and cause cardiomyopathy. However, measurements of the intrinsic force, velocity and ATPase activityof myosin have not provided a consistent mechanism to link mutations to muscle pathology. An alternative modelpositsthat mutations in myosin affect the stability ofa sequestered, super-relaxed state (SRX) of the proteinwith very slow ATP hydrolysis and thereby change the number of myosin heads accessible to actin. Here, using a combination of biochemical and structural approaches, we show that purified myosin enters aSRX thatcorresponds to a folded-back conformation, which in muscle fibersresults insequestration of heads around the thick filament backbone. Mutations that cause HCM destabilize this state, while the small molecule mavacamtenpromotes it. These findings provide a biochemical and structural link between the genetics and physiology ofcardiomyopathywith implications for therapeutic strategies.

## Introduction

Muscle myosin is a hexamer consisting of two myosin heavy chains and two sets of light chains, the essential light chain (ELC) and the regulatory light chain (RLC). The myosin molecule can be divided into two parts, heavy meromyosin (HMM),which consists of two globular heads and the first ∼40% of the coiled-coil tail, and light meromyosin (LMM),which consists of the C-terminal ∼60% of the coiled-coil tail. LMM self assembles creating the shaft of the myosin thick filament found in sarcomeres of the muscle. HMM can be further divided into Subfragment 1 (S1), which is the globular head of the myosin that serves as the motor domain (Toyoshima et al., 1987),and subfragment 2 (S2). S1 houses the ATP and actin binding sites followed by anessential light chain (ELC) and regulatory light chain (RLC)bound α-helix (lever arm) (Fig.1A). S1 heads are arranged on the thick filament backbone in muscle in a helical or quasi-helical fashion. There is a 14.3-nm vertical spacing between two adjacent myosin molecules on the filament with a true repeat of 42.9 nm.

**Fig. 1.**
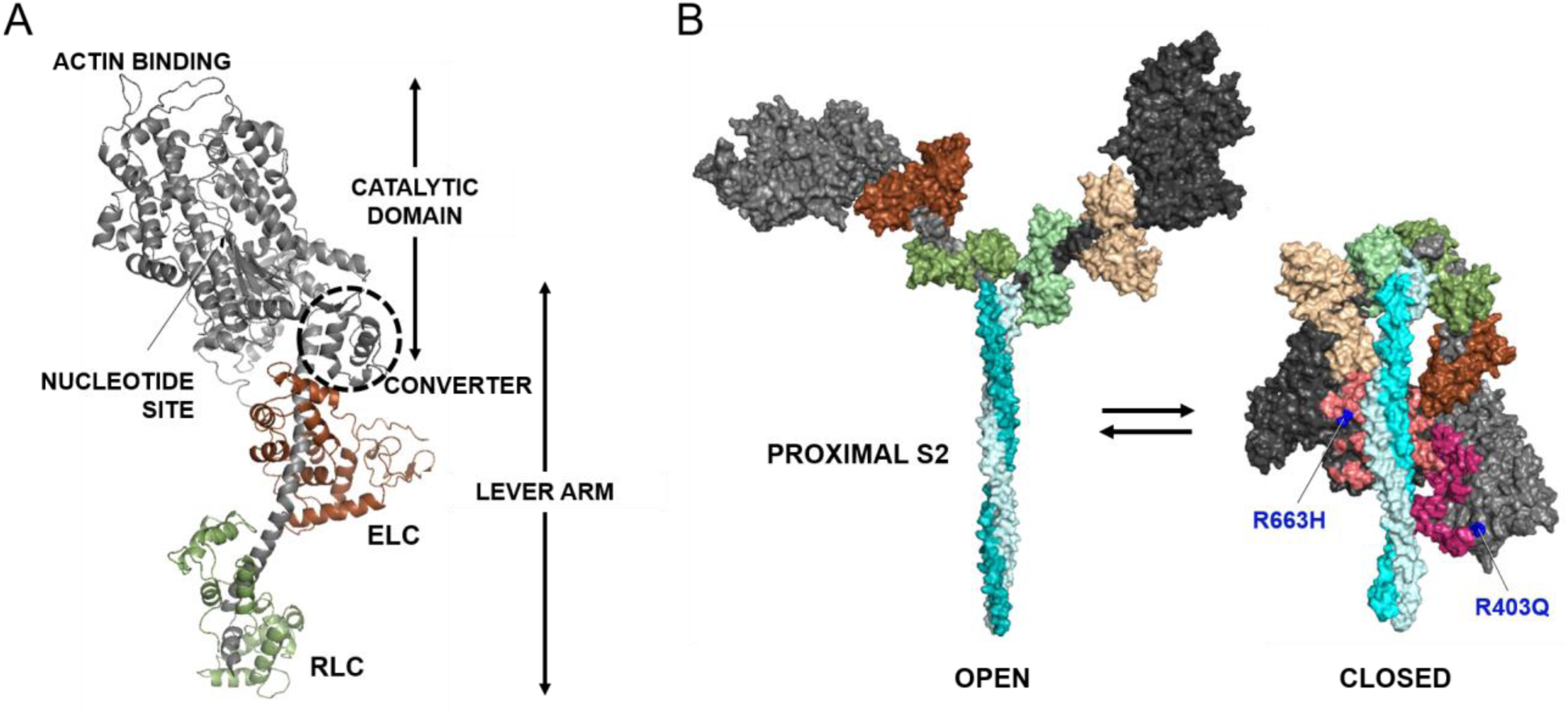
Human β-cardiac myosin structural models. **(A)** Pymol homology model of human β-cardiac myosin subfragment 1 (S1) in the pre-stroke state showing its various domains. ELC,essential light chain (brown); RLC, regulatory light chain (green). Modeling was done as described previously (Nag et al., 2017; Trivedi et al., 2017). **(B)**Structural models of the open on-state and the closed IHM off-state of human β-cardiac myosin. The motor domains are black (blocked head) and dark gray (free head), while the ELCs are in shades of brown, and RLCs in shades of green. The positions of R403Q and R663H (blue) are shown in the IHM relative to the mesa residues (Trivedi et al., 2017) (light pink, blocked head; dark pink, free head). Modeling was done as described previously (Nag et al., 2017; Trivedi et al., 2017).

Importantly, intramolecular interactions favoring a folded state of myosin (Fig. 1B) have been observed in both purified myosin preparations(Burgess et al., 2007; Jung et al., 2011; Jung et al., 2008; Wendt et al., 1999; Wendt et al., 2001) and in myosin thick filaments isolated from muscle (Al-Khayat et al., 2013; Alamo et al., 2008; Woodhead et al., 2005; Zoghbi et al., 2008). Named the interacting-heads motif (IHM) (Alamo et al., 2008), this structure has been proposed to be related to a super-relaxed state (SRX) of muscle, defined as a state that has a much reduced basal ATPase rate of ∼ 0.003 s^-1^(McNamara et al., 2015). The SRX was initially discovered in rabbit skeletal muscle and thereafter also seen in rabbit cardiac, tarantula exo-skeletal and mouse and human cardiac fibers (Hooijman et al., 2011; McNamara et al., 2016; Naber et al., 2011; Stewart et al., 2010). An appealing hypothesis is that the SRX state is related to theIHM myosin state where the myosin S1 heads are interacting with one another and folded back onto their own coiled-coil S2 tail (Alamo et al., 2016; Cooke, 2011; Hooijman et al., 2011; Nogara et al., 2016; Wilson et al., 2014). Since the SRX state has only been described in fibers, it is possible that other sarcomeric proteins in the vicinity of the myosin are essential for establishing this state. There have been no studies that directly demonstrate a structural corollary of the SRX state with purified sysems.Here we demonstrate biochemically that a decrease of basal ATPase to the levels seen in the SRX in fibers can be observed with purified human β-cardiac myosin alone.

Hypertrophic cardiomyopathy (HCM) is an autosomal dominant inherited disease of heart muscle(Geisterfer-Lowrance et al., 1990; Seidman and Seidman, 2000; Seidman and Seidman, 2001)characterized by hyper-contractility and subsequent hypertrophy of the ventricular walls. Systolic performance of the heart is preserved or even increased, but relaxation capacity is diminished. Patients with HCM are at increased risk of heart failure, atrial fibrillation and stroke, and sudden cardiac death.Current pharmacological management options are not disease specific and do not address the underlying HCM disease mechanism. It has been hypothesized that a small molecule that binds directly to myosin and normalizes the hyper-contractility of the sarcomere may interrupt the development of downstream pathology (such as hypertrophy, fibrosis and clinical complications) seen in this disease (Spudich, 2014). Mavacamten (formerly MYK-461; MyoKardia Inc., South San Francisco, CA) is anoral allosteric modulator of cardiac myosin and causes dose-dependent reductions in left ventricular contractility in healthy volunteers and HCM patients(Maron et al., 2016). This investigational drug is in phase 2 clinical trials and is the first in line of potential new HCM therapeutic agents that act directly by interacting with the human β-cardiac myosin and normalizing its power output. Mavacamtenwas identified in a screen for actin-activated myosin ATPase inhibitors, and it lengthens the total cycle time (t_c_) of the ATPase cycle(Green et al., 2016). This results in a reduction of the duty ratio (t_s_/t_c_) and therefore of the ensemble force, since F_ens_ = f_int_· N_a_· duty ratio, where f_int_ is the intrinsic force of a myosin molecule and N_a_ is the total number of myosin heads in the sarcomere that are functionally accessible for interaction with actin. Here we show that mavacamtenalso reduces N_a_ by stabilizing the heads in a folded IHM state with a very slow release of bound nucleotide, an SRX-IHM state.

The first HCM-causing mutation in β-cardiac myosin to be identified was R403Q (Geisterfer-Lowrance et al., 1990). As with other HCM mutations, the R403Q mutation results in hyper-contractility of the muscle(Tyska et al., 2000; Wilson et al., 1967). We provide evidence here that the R403Q mutation results in a shift in the equilibrium between the open and closed structures of myosin (Fig. 1B) to the open state, resulting in more heads available for interaction with actin and the hyper-contractility seen clinically. We see a similar phenotype in human cardiac samples with the HCM-causing R663H mutation. We further demonstrate that mavacamten reverses this destabilization of IHM caused by these mutations.

## Results

### Purified human ß-cardiac myosincontaining the proximal S2 portion of its coiled-coil tail is in an ionic-strength dependentsuper-relaxed state

To test the hypothesis that the SRX observed in muscle fibers is related to the folded-back IHM of purified myosin we compared the kinetics of myosin’s basal ATPase cycle for three purified human ß-cardiac myosin constructs: 25-hep HMM (two-headed with first 25-heptads of proximal S2), 2-hep HMM (two-headed with first 2-heptads of proximal S2) and short S1 (sS1; single-headed with no S2, and truncated immediately after the ELC binding domain, Fig. 1A). The basal ATPase rates of the three constructs were all within the range of 0.01 to 0.03 s^-1^ (Fig. S1), consistent with high levels of actin-activation of these human ß-cardiac myosin constructs (about 100-fold activation by actin(Adhikari et al., 2016; Nag et al., 2017). SRX is defined as a rate of nucleotide release from the myosin head that is even slower, about 0.003 s^-1^(McNamara et al., 2015).

To test the percentage of heads in the SRX for the three constructs weadapted the single nucleotide-turnover assay, which is typically measured in a stopped-flow apparatus, to a plate-based measurement. The slow rates of normal myosin basal ATPase and the SRX-based nucleotide release makes them amenable to a plate-based measurement. Moreover, owing to the difficulties in expressing high amounts of human cardiac proteins, the plate-based assay is an apt choice for this kind of experiment. This assay measured the fluorescentnucleotide release rates by loading the myosin heads with mant-ATP and then chasing with excess unlabeled ATP, as had been done in skinned fibers previously (McNamara et al., 2015). As the mant-nucleotide is released from the myosin, its fluorescence decreases(McNamara et al., 2015).

The decay rate of mant-nucleotide fluorescence for 25-hep HMM was fit well by two exponential rate constants, one representing a basal ATPase rate of heads in a presumed open state (disordered relaxed state, DRX) of ∼0.03 s^-1^and the other representing an SRX rate of ∼0.003 s^-1^ (Fig. 2A). In 100 mM potassium acetate (KAc), the amplitudes were 74 ± 2% basal rate and 26 ± 2% SRX rate (Fig. 2B). This SRX level observed for the 25-hep HMM did not vary significantly from preparation to preparation, and may largely represent myosin heads that are folded back onto their own proximal S2 tail (Fig. 2C). Consistent with this hypothesis, the fraction of myosin heads in the SRX increased from 26 ± 2% to 42 ± 2% (p<0.05) when the KAc was decreased from 100 mM to 25 mM, and further increased from 26 ± 2% to 59 ± 7% (p<0.05) when the salt was decreased from 100 mM to 5 mM KAc (Fig. 2B). This would be expected since the IHM structure is thought to be held together largely by charge-charge interactions (Alamo et al., 2017; Alamo et al., 2008; Blankenfeldt et al., 2006; Moore et al., 2012; Nag et al., 2017). The amplitudes of the fast (74 ± 2%) and slow phases (26 ± 2%)observed with the human ß-cardiac 25-hep HMM at 100 mM KAcmatch closely with the amplitudes observed in SRX experiments done with rabbit cardiac fibers (∼70% fast and ∼30% slow at 120mM KAc)(Hooijman et al., 2011).

**Fig. 2.**
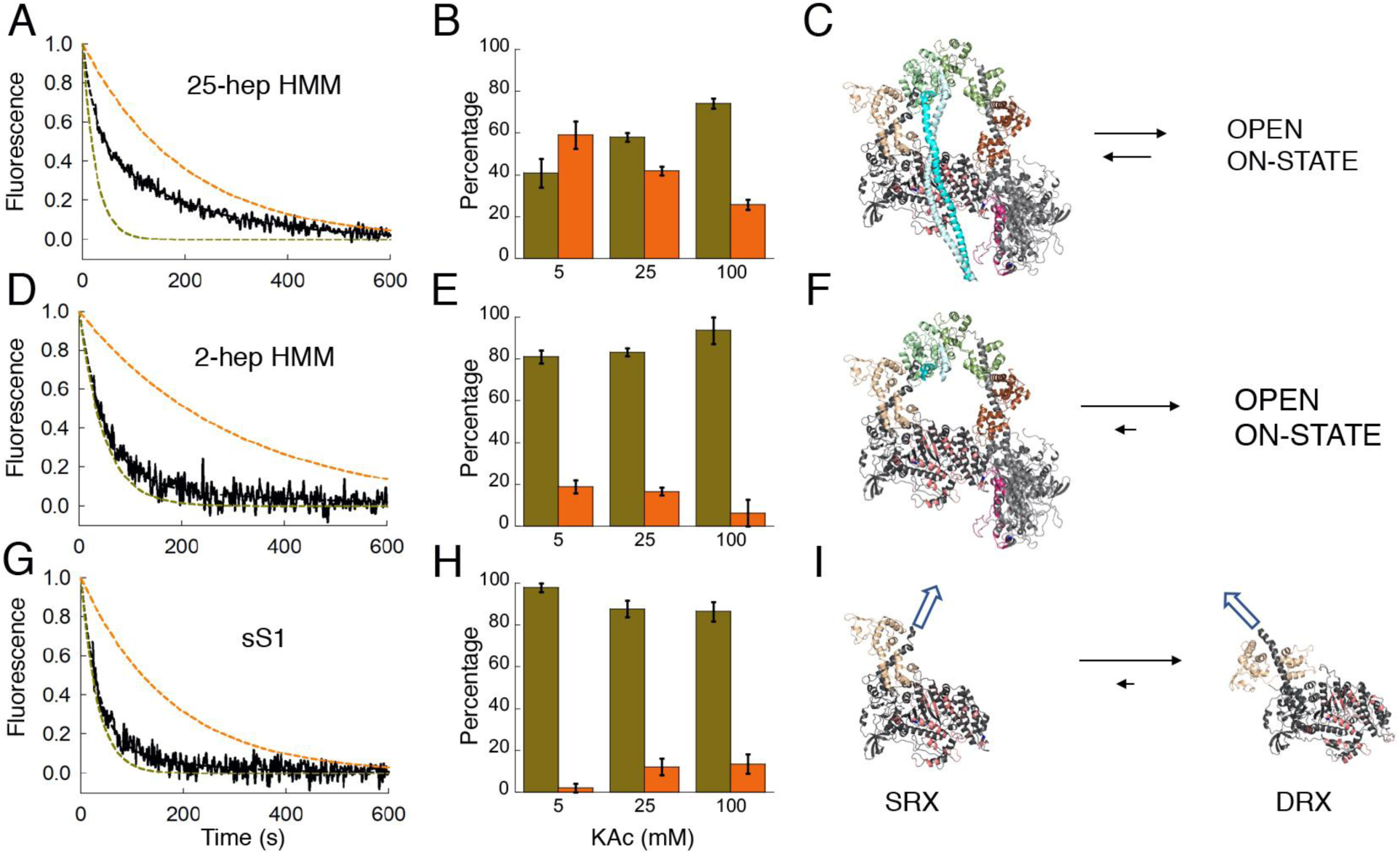
Mant-nucleotide release rates for purified human ß-cardiac myosin fragments. **(A, D, G)** Fluorescence decays ofmant-nucleotide release from 25-hep HMM, 2-hep HMM and sS1, respectively, in 25 mM KAc (solid black curves). The simulated single-exponential orange dashed curvesare the SRX rate (0.003 - 0.005 s^-1^) and the simulated single-exponential dashed olive green curvesarethe DRX rate (0.02 – 0.04 s^-1^). The simulated orange and olive green dashed curves act as references for the single exponential fits of slow and fast phases, respectively, that derive from fitting the solid black data curves with the best two exponential fits. Thus, the data (black lines) were fit with a combination of these two single exponentials. **(B, E, H)**Percentage of myosin heads in the SRX (orange) versus DRX (olive green)statescalculated from the amplitudes of the double-exponential fits of the fluorescence decays corresponding to the mant-nucleotide release from the 25-hep HMM, 2-hep HMM and sS1 constructsat various KAc concentrations. **(C)**Homology model of folded-backhuman ß-cardiac 25-hep HMM (MS03, https://spudlab.stanford.edu/homology-models/, showing only 13 heptad repeats of the S2) in equilibrium with an open on-state;**(F)** 2-hep HMM (from MS03) in equilibrium with an open on-state; and**(I)** sS1 SRX (left, the homology-modeled blocked-head only from MS03) in equilibrium with sS1 DRX (right, from HBCprestrokeS1; the modeled relaxed ATP and/or ADP.Pi bound states of the human ß-cardiac myosin heads). The white arrows illustrate the different directions the light-chain binding regions point in the SRX vs DRX states, where all alignments in (C, F, and I) are the same. Error bars denote s.e.m.

Interestingly, the 2-hep HMM, which contains only 2 heptad repeats of the S2 tail, still showed19± 3%SRXin 5 mM KAc(Fig. 2D,E), but the fractions of SRX at different KAc concentrations were not statistically significant(p > 0.05). It is possible that some blocked head interactions with the free head help stabilize this level of SRX in the 2-hep HMM (Fig. 2F). However, our finding of a similar level of SRX (∼15%) in our sS1 preparations (Fig. 2G,H) suggests an alternative explanation. It is important to note that myosin molecules in solution in any nucleotide state are likely to exist in a number of closely-related conformational states in equilibrium. We think it possible, therefore, that ∼15% of the population of independent sS1 molecules in solution exist in a conformation that is similar to that stabilized in the IHM structure (Fig. 2I, the SRX) in equilibrium with DRX heads (pre-stroke heads having ATP or ADP.Pi bound). In the context of the 25-hep HMM, this conformation favors the interactions with the proximal S2 and other head-head interactions that define the SRX-IHM. Thus, strictly speaking, myosin heads can occupy the SRX without entering the IHM state directly, but the SRX conformation of myosin heads isfurther stabilized by intramolecular interactions in the SRX-IHM. This idea gains strength by the mavacamten experiments described below.

To ensure that we were not missing any fast phase in our plate-based measurement, we performed the similar measurement with sS1 in a stopped-flow apparatus. The rates and amplitudes obtained were very similar to our plate-based measurements (Fig. S2).

These results suggest that purified myosin can adopt a biochemical SRXthat is stabilized by the S2 tail and by low ionic strength conditions that favor electrostatic interactions. These are the first results that directly link, using purified human ß-cardiac myosin, the IHM structural state with the biochemical SRX.

### The cardiac myosin inhibitor mavacamten stabilizes the SRX-IHMof purified human ß-cardiac myosin

Using the mant-nucleotide release rates as a measure of the level of SRX in the population, we examined whether the cardiac myosin inhibitor mavacamten has any effect on the distribution of ATPase states in the population of purified human ß-cardiac myosin constructs. Mavacamten was selected on the basis that it inhibits the actin-activated activity of sS1 (Green et al., 2016), but the mechanism of that inhibition is unknown. Here we show that it has a profound effect on stabilizing the SRX. Mavacamten induced a 20-fold reduction in the basal ATPase rate of the 25-hep HMM (Fig. S1A). In the single-nucleotide turnover assay, in the presence of 10 µM mavacamten, the mant-nucleotide release rates from 25-hep HMM at 25 mM KAcwere fit well by a single exponential with a rate constant reflecting the SRX. At both 25 mM KAc (Fig 3A,B) and 100 mM KAc (Fig. S3), essentially ∼100% of the myosin heads were in the SRX in the presence of mavacamten. In absence of mavacamten, the fraction of 25-hep HMM in the SRX was ∼42% at 25 mM KAc (Fig. 2B, Fig. 3B) and ∼26% at 100 mM KAc (Fig 2B).

**Fig. 3.**
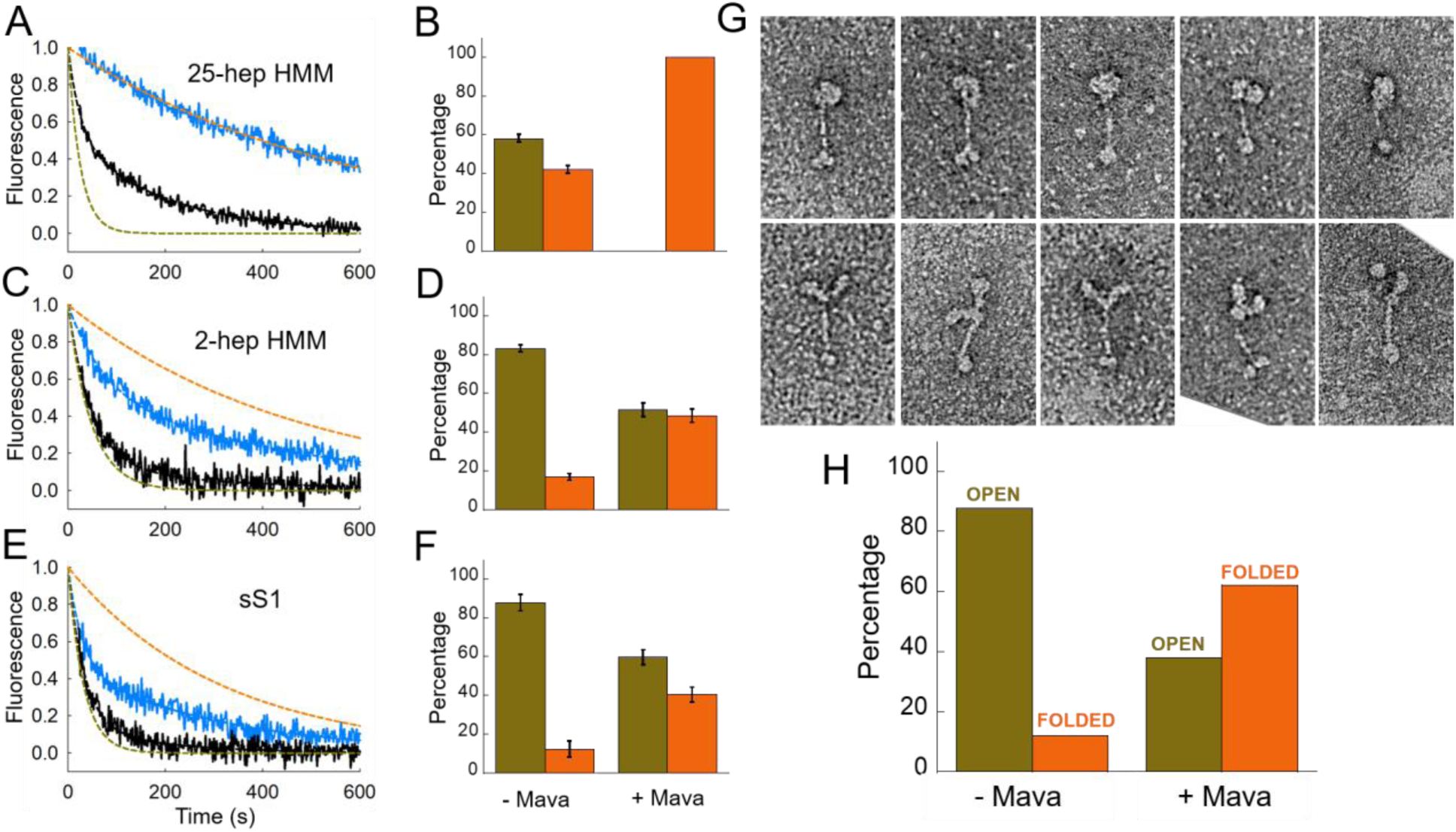
Mant-nucleotide release rates and percentage folded states for purified human ß-cardiac myosin fragments with and without mavacamten. **(A, C, E)** Fluorescence decays of mant-nucleotide release from 25-hep HMM, 2-hep HMM and sS1 in 25 mM KAc with (blue curve) and without (black curve) mavacamten. The simulated orange and olive green dashed curves act as references for the single exponential fits of slow and fast phases, respectively, that derive from fitting the solid black data curves with the best two exponential fits. **(B, D, F)** Percentage of myosin heads in the SRX (orange) versus DRX (olive green)stateswith and without mavacamten calculated from the double-exponential fits of the fluorescence decays corresponding to the mant-nucleotide release from 25-hep HMM, 2-hep HMM and sS1 at 25 mM KAc.These data derive from three independent protein preparations each of 25-hep HMM, 2-hep HMM and sS1 made on different days. **(G)** Panel showing representative single particle negatively-stained electron microscopy images of the open and closed forms of the 25-hep HMM. Both the open and closed molecules have been selected from a sample containing mavacamten crosslinked to 25-hep HMM at 25 mM KAc. **(H)** Percentage of 25-hep HMM molecules in the closed or open form in the presence and absence of mavacamten. Total of 622 and 183 molecules were counted for the with and without mavacamten samples, respectively. Error bars denote s.e.m.

The effect of mavacamten on 2-hep HMM was less pronounced (Fig. 3C,D), consistent with a role for the proximal S2 in stabilizing the SRX. However, 40-50% SRX was observed in the presence of mavacamten suggesting that either the heads can interact with one another to some extent in the absence of proximal S2 and this is stabilized by mavacamten binding (shifting the equilibrium to the left in Fig. 2F) or mavacamtenstabilizes the SRX conformation of sS1 in equilibrium with DRX heads (shifting the equilibrium to the left in Fig. 2I).

Indeed, mavacamten increases the SRX levels of sS1 to nearly that of the 2-hep HMM (Fig.3F vs D). Mavacamten increased the SRX levels of 2-hep HMM to 48 ± 4% and sS1 to 40 ± 4%, which are not significantly different (p>0.05). This is also reflected in a reduction of the basal myosin ATPase of the 2-hep HMM and sS1 constructs by the drug (Fig. S1B, C). Thus, we think it likely that the binding of mavacamten to sS1 alone stabilizes an SRX off-state, which is favored to fold into the IHM structure (Fig. S4).We hypothesize that the sS1 SRX conformation will have its light-chain binding helix tilted a bit further in the pre-stroke direction than normally achieved by the relaxed DRX pre-stroke state (Fig. S4). Since the human ß-cardiac myosin homology model is derived from a low-resolution (2 nm) tarantula skeletal muscle template, a mava-bound sS1 structure may represent a more realistic and higher-resolution version of the blocked head of the SRX-IHM, and possibly relate to the free head of the SRX-IHM as well.

We used electron microscopy to verify that the SRX observed with 25-hep HMM corresponds to folded-back (closed) heads. Fig. 3G shows representative images of closed IHM structures (top panels) and open structures (lower panels). After coupling the 25-hep HMM to a mavacamten analog containing a photoactivatable crosslinker, folded-back IHM structures dominated the images, while the reverse was true in the absence of the mavacamten derivative (DMSO control)(Fig. 3H). Less than 20% of the myosin molecules were in a closed IHM state in the absence of mavacamten and ∼60% were in a closed IHM state in the presence of mavacamten (Fig. 3H). These results more firmly link the IHM structural state with the SRX.

### The cardiac myosin inhibitor mavacamten stabilizes the SRX-IHMof porcine and human cardiac muscle fibers

To examine whether the effects of mavacamten on purified myosin translate to an increase in the level of SRX in cardiac fibers, we carried out mant-nucleotide release analyses on skinned fibers from relaxed minipigcardiac muscle. Mavacamten increased the percentage of the SRX in the porcine fibers from 26 ± 2% to 39 ± 4% (p<0.05, Fig. 4A,B). There is a corresponding decrease in tension measured in skinned porcine cardiac muscle fibers exposed to mavacamten from 14.4 ± 0.7 mN/mm^2^ to 6.6 ± 0.7 mN/mm^2^ (pCa5.8, p < 0.05, Fig 4C).

**Fig. 4.**
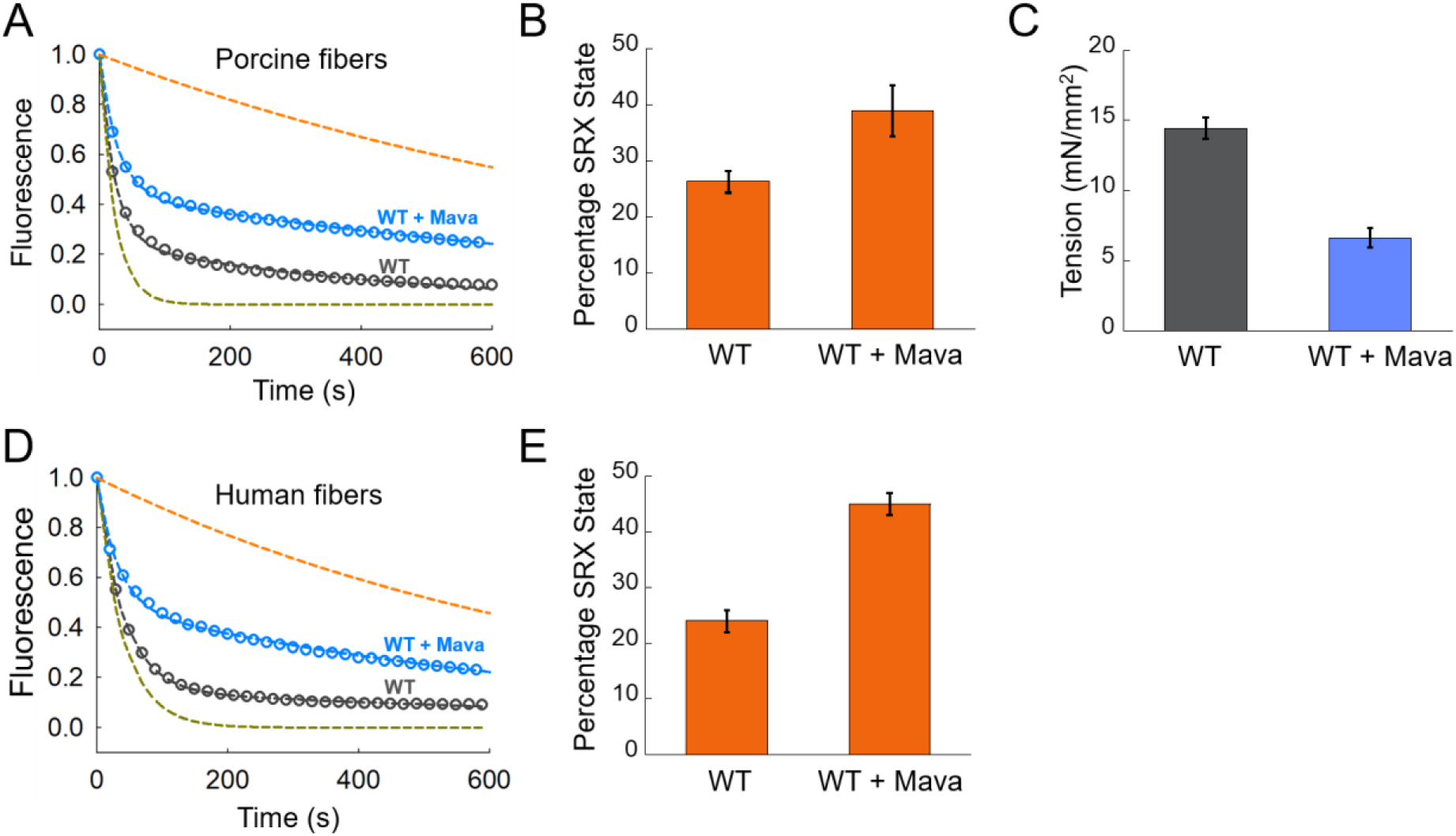
Effect of mavacamten on wild-type porcine and human cardiac fibers. Blue and dark gray denote experimental conditions with and without mavacamten, respectively. **(A)** SRX measurements with skinned cardiac fibers from wild-type pigs in the presence and absence of mavacamten. The simulated orange and green dashed curves act as references for the single exponential fits of slow and fast phases, respectively, that derive from fitting the solid blue and black data curves with the best two exponential fits. **(B)** Percentage of SRX detected in the porcine cardiac fibers with and without mavacamten. **(C)** Maximum tension measurement in the skinned porcine cardiac fibers in the presence and absence of mavacamten. **(D)** SRX measurements with skinned cardiac fibers from human hearts in the presence and absence of mavacamten**. (E)** The percentage of SRX in human cardiac fibers in the presence and absence of mavacamten. Error bars denote s.e.m.

A similar mavacamten-mediated SRX stabilization effect was also observed for wild type (WT) human cardiac fibers. Mavacamten increased the SRX in the human fibers from 24 ± 2% to 45 ± 2% (p<0.05, Fig. 4D,E). The smaller effect on the fibers compared to the purified 25-hep protein could reflect a role for other components of the intact sarcomere in regulating the SRX population, or accessibility issues of mavacamten to the myosin in the skinned fibers. The ∼25% SRX we see in the fibers without mavacamten treatment at 120 mM KAc is consistent with our 25-hep HMM purified protein data at 100 mM KAc (Fig. S3B).

To examine the spatial organization of the myosin heads weused low-angle X-ray diffraction of porcine cardiac muscle fibers, which reveals information about the degree of order of the myosin heads along the myosin thick filaments as well as their proximity to the thin actin filaments. A representative low-angle X-ray diffraction image of a porcine cardiac muscle fiber shows the typical helical diffraction pattern known for striated muscle (Fig. 5).

**Fig. 5.**
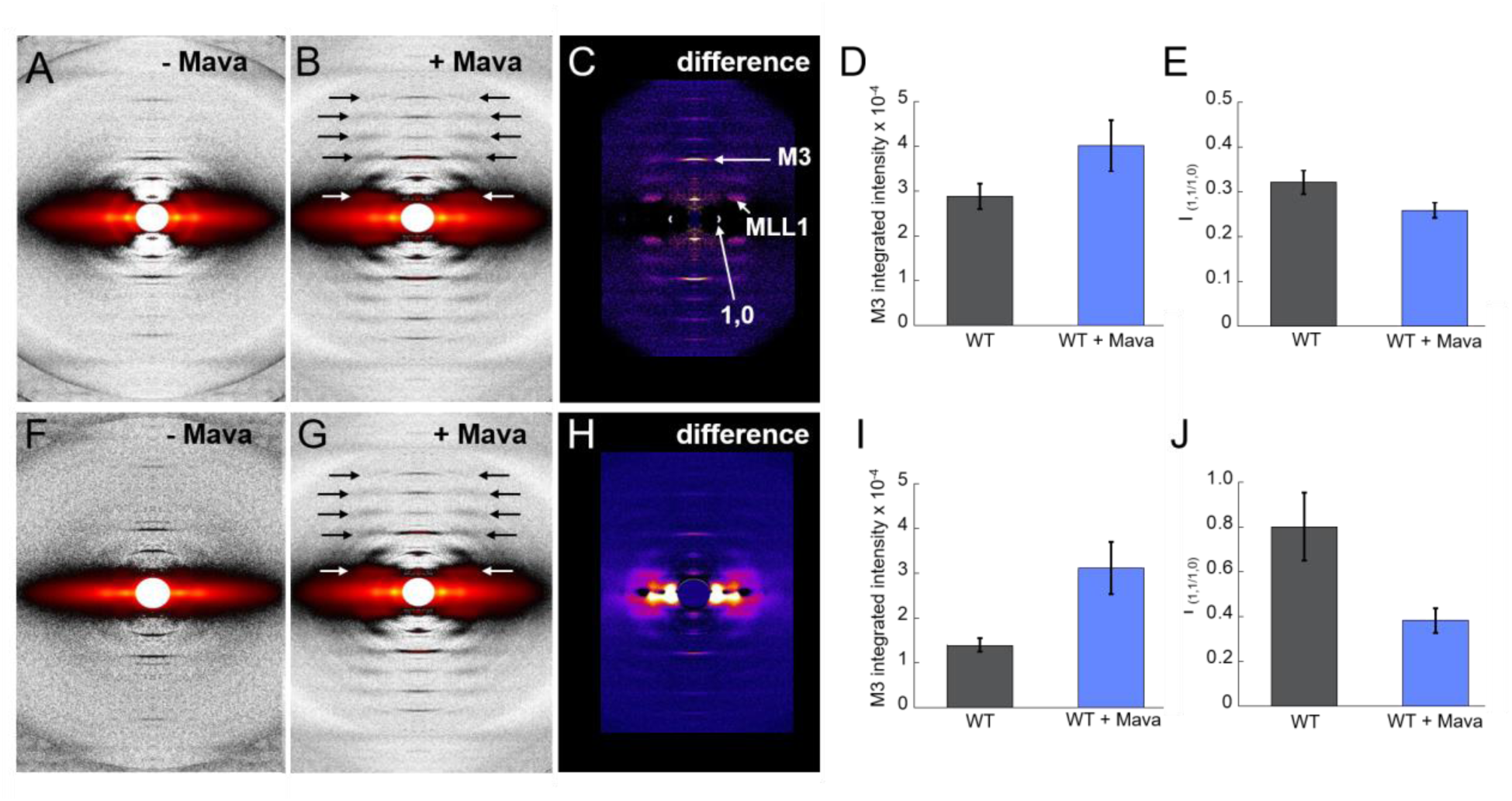
Effect of mavacamten on ordering of myosin heads onto the thick filaments in wild-type porcine fibers. (A, B, C) Diffraction patterns of wild-type porcine fibers under relaxing conditions (pCa 8) without and with mavacamten and the difference in intensities. **(D)** Mavacamten increases the intensity of the meridional reflection M3 under relaxing conditions. **(E)** Mavacamten decreases the equatorial intensity ratio, I_1,1/1,0_, under relaxing conditions. **(F, G, H)** Diffraction patterns of wild-type porcine fibers under contracting conditions (pCa 4) without and with mavacamten and the difference in intensities. **(I)**Mavacamten increases the intensity of the meridional reflection M3 under contracting conditions. **(J)** Mavacamten decreases the equatorial intensity ratio, I_1,1/1,0_,under contracting conditions. In (D), (E), (I) and (J), blue and dark gray bars denote experimental conditions with and without mavacamten, respectively. Error bars denote s.e.m.

Mavacamten causes a dramatic ordering of myosin heads along the backbone of the thick filaments as observed by the substantial increase in the myosin-based helical layer line reflections for both relaxed (Fig. 5A vs B, arrows) and contracting (Fig. 5F vs G, arrows) porcine muscle fibers. The first myosin layerline (MLL1) intensity at 43 nm (Fig. 5B,G white arrows) increased about 40% with mavacamten treatment (MLL1_+mava_/MLL1_-mava_= 1.37±0.13). If we assume that only myosin heads in the SRX give rise to layer lines, a 40% MLL1 intensity increasewould imply 17±6% more heads in the SRX, consistent with the fluorescence decay data shown in Fig. 4A,B. A prominent reflection is the M3 reflection on the meridian, which reports on the 14.3-nm repeat of the myosin heads along the longitudinal shaft of the thick filamentin the relaxed state(Fig. 5C). The higher the intensity of this reflection, the more ordered are the heads along this 14.3-nm repeat. When the diffraction pattern of the relaxed muscle in the absence of mavacamten (Fig. 5A) is subtracted from the diffraction pattern in the presence of mavacamten (Fig. 5B), the difference (Fig. 5C) shows that the intensity of the M3 reflection is significantlyincreased by addition of mavacamten (p < 0.05). This is quantified in Fig. 5D where the results from 13fibers are shown. An even more dramatic increase is seen in the M3 reflection when mavacamten is added to contracting fibers (p < 0.05, Fig. 5F-I). Thus, for both relaxed and contracting fibers, mavacamtenincreases the helical order of myosin heads on the thick filament backbone.

The 1,1 and 1,0 reflections on the equator provide information on the number of myosin heads that have moved away from the myosin thick filaments toward the actin thin filaments. The 1,1 lattice planes include the actin filament densities, while the 1,0 lattice planes include only the myosin thick filament densities. Thus an increase in the intensity of the 1,0 reflection shows that more myosin heads have been sequestered near the shaft of the thick filaments, and the I_1,1_/I_1,0_ intensity ratio is reduced. As can be seen in Fig. 5C and H, mavacamten causes an increase in the 1,0 reflection intensity. Furthermore, Fig. 5E and J show that the I_1,1_/I_1,0_ intensity ratio is reduced (p < 0.05) by mavacamten in both relaxing and contracting conditions, suggesting that it causes more heads to become sequestered against the LMM backbone of the myosin thick filament.

These structural results, combined with the studies with purified human ß-cardiac myosin described above, suggest that mavacamten shifts the open-closed state equilibrium toward the closed SRX-IHM state, which in muscle fibers is more ordered near and along the backbone of the myosin thick filament.

### The hypertrophic cardiomyopathy mutations R403Q and R663H reduce the level of SRX in cardiac muscle fibers

To better characterize the effects of mutations that cause HCM on cardiac structure and function, we generated a large animal model of HCM in the Yucatan minipig (Fig. S5 and Methods). We introduced into minipigs the heterozygous *MYH7* R403Q mutation, which has been well-characterized in mice and humans (Geisterfer-Lowrance et al., 1996; Geisterfer-Lowrance et al., 1990). When cardiac fibers from the R403Q pigs were compared with those from the WT pigs for SRX levels, strikingly, the R403Q mutation caused a significant decrease in the percentage of SRX in the porcine fibers, from 26 ± 2% to 20 ± 2% (p<0.05, compare Fig. 6A,B with Fig.4A,B). This result supports the hypothesis that myosin HCM mutations cause clinical hyper-contractility by shifting the equilibrium fromthe closed off-state (SRX-IHM) to the open on-state where the heads are free to interact with actin (Alamo et al., 2017; Nag et al., 2017; Nag et al., 2016; Spudich, 2015; Trivedi et al., 2017). Addition of mavacamten to the R403Q fibers returned the percentage of SRX back to the normal WT level (30 ± 5%, p<0.05) and also reduced tension from 14.9 ± 0.7 mN/mm^2^ to 8.1 ± 0.6 mN/mm^2^ (pCa5.8, p < 0.05, Fig. 6C).The physiologically-relevant pCa range for tension development in the heart is from ∼10^-7^ M Ca^2+^ to ∼10^-6^M Ca^2+^ (redshaded area, Fig. S6)(Previs et al., 2015), where the R403Q mutation induces hyper-contractility and mavacamten reduces it towards normal (light blue to dark blue shaded area, Fig. S6).

**Fig. 6.**
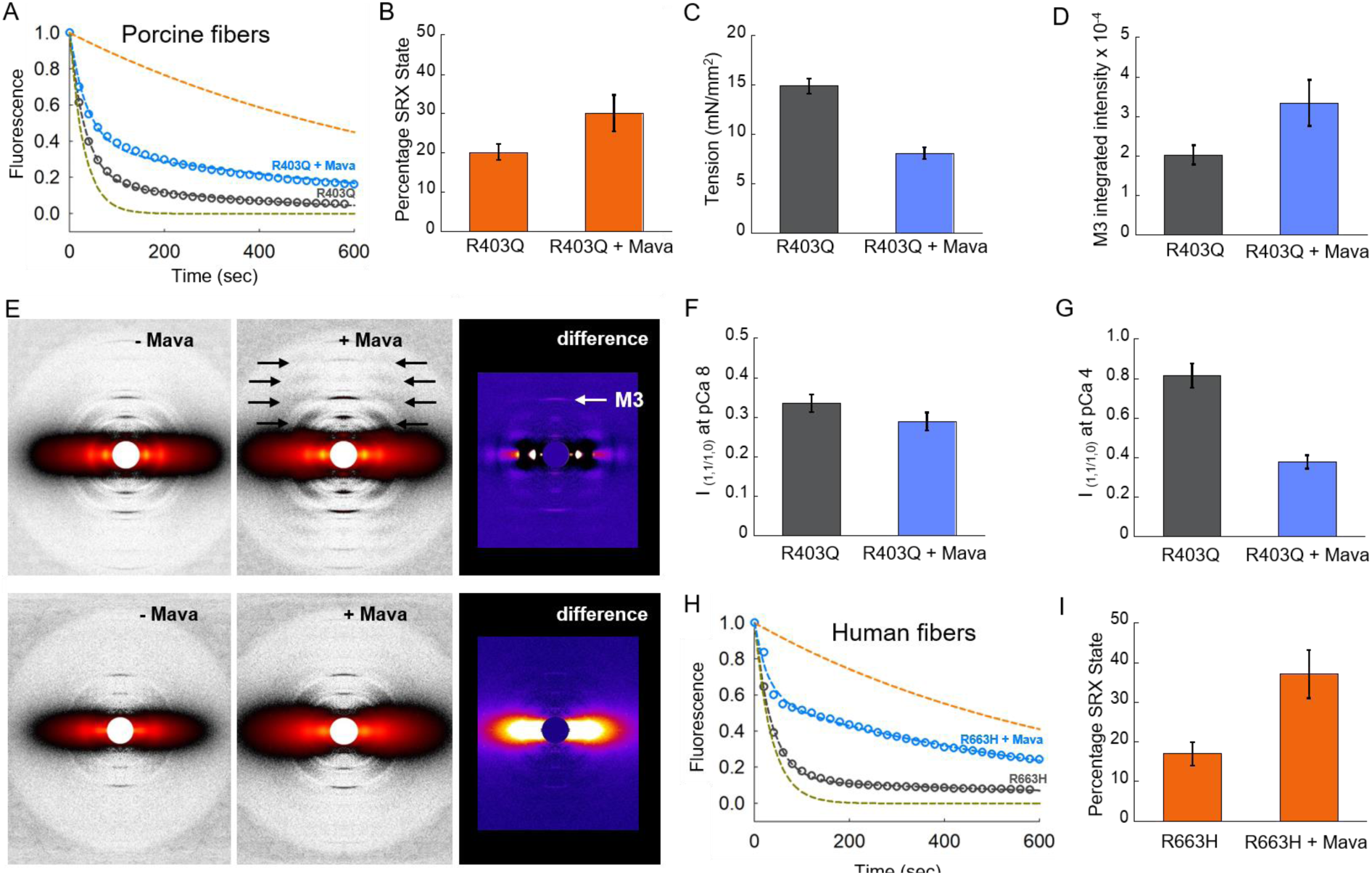
Mavacamten stabilizes the SRX in porcine and human cardiac fibers carrying HCM mutations. Blue and dark gray denote experimental conditions with and without mavacamten, respectively. **(A)** SRX measurement with skinned cardiac fibers from R403Q pigs in the presence and absence of mavacamten. The green and orange curves are simulated single-exponential fits of fast and slow phase, respectively. **(B)** Percentage of SRX detected in the porcine cardiac fibers with and without mavacamten. **(C)** Maximum tension measurement in the skinned cardiac fibers from R403Q pigs in the presence and absence of mavacamten. **(D)** Mavacamten increases the intensity of the meridional reflection M3 under relaxing conditions. **(E)** Diffraction patterns of R403Q porcine fibers under relaxing conditions (pCa 8) without and with mavacamten and the difference in intensities (top panel). Diffraction patterns of R403Q porcine fibers under contracting conditions (pCa 4) without and with mavacamten and the difference in intensities (bottom panel). **(F)** Mavacamten decreases the equatorial intensity ratio, I_1,1/1,0_, of R403Q porcine fibers under relaxing conditions. **(G)** Mavacamten decreases the equatorial intensity ratio, I_1,1/1,0_, of R403Q porcine fibers under contracting conditions. **(H)** SRX measurements with skinned cardiac fibers from R663H human hearts in the presence and absence of mavacamten. **(I)** The percentage of SRX with and without mavacamten in the R663H human cardiac fibers. Error bars denote s.e.m.

Treatment of R403Q skinned porcine fibers with mavacamtencaused an increase in ordering of myosin heads along the backbone of the myosin thick filament, as judged by an increase in the myosin-based helical layer lines (black arrows) and the intensity of the M3 reflection(white arrow) (Fig. 6E), consistent with a shift in equilibrium toward the myosin SRX-IHM off-state (Fig. 6D,E). This mavacamten-dependent increase in the intensity of the M3 reflection was observed under relaxing conditions. Such an intensity change was difficult to quantify for the R403Q fibers under contracting conditions. Mavacamtenalso caused asignificant decrease in the I_1,1_/I_1,0_ intensity ratio of the R403Q fibers (p < 0.05) (Fig. 6E-G), further supporting a shift in equilibrium toward the myosin SRX-IHM off-state more closely associated with the backbone of the myosin thick filament. This effect was quantified for the R403Q fibers under both relaxing and contracting conditions.

We also studied the properties of the SRX in a known human pathogenic HCM mutation, *MYH7* R663H with fibers isolated from human cardiac tissue. We found that the R663H mutation caused a significant decrease in the percentage of SRX from 24 ± 2% (WT human cardiac tissue) to 18 ± 2% (R663H human cardiac tissue) (p<0.05), and that the addition of mavacamten to these fibers restored the SRX levels back to 37 ± 6% (p<0.05), which is similar to the WT level (Fig. 6H,I). These changes are similar to those observed with the R403Q porcine samples and suggest that HCM mutations in myosin destabilize SRX in both pigs and humans.

### Discussion

Over the past five decades our understanding of muscle physiology, biochemistry and biophysics has greatly advanced. While not complete, we have a good understanding of how many of the most important sarcomeric proteins function and their roles in muscle contractility. Now focus is shifting to understanding the molecular basis of disease states of these proteins and modern therapeutic approaches that can be applied to them. This study focuses on HCM mutations that cause hyper-contractility of the heart and a small molecule, mavacamten, which resets this hyper-contractility to normal.

Recent studies using human ß-cardiac myosin support the view that many, if not most, myosin missense HCM mutations cause the hyper-contractility observed clinically by shifting an equilibrium between a sequestered off-state of myosin heads to their on-state now able to interact with actin (Adhikari et al., 2016; Alamo et al., 2017; Homburger et al., 2016; Kawana et al., 2017; Nag et al., 2017; Spudich, 2015; Spudich et al., 2016; Trivedi et al., 2017). The majority of these mutations are highly localized to three surfaces of the myosin molecule: the converter domain (Alamo et al., 2017; Homburger et al., 2016) (Fig. 1A), a relatively flat surface named the myosin mesa (Spudich, 2015) (Fig. 1B), and the proximal part of S2 that interacts with the S1 heads in the IHM (Adhikari et al., 2016; Alamo et al., 2017; Homburger et al., 2016; Kawana et al., 2017; Nag et al., 2017; Spudich et al., 2016; Trivedi et al., 2017) (Fig. 1B).

The realization that the myosin mesa of the two S1 heads in the IHM are in fact cradling the proximal S2 (Fig. 1B) was striking(Adhikari et al., 2016; Nag et al., 2017). Moreover, the mesa is highly enriched in HCM mutations in positively-charged Arg residues(Spudich, 2015), whereas the proximal S2 is highly enriched in HCM mutations in negatively-charged Glu and Asp residues (Homburger et al., 2016).Furthermore, the converter HCM mutations are all in a position to potentially disrupt the IHM at a primary head-head interaction site (PHHIS) involving a particular surface of the ‘blocked head’ of the IHM and the converter of the ‘free head‘(Alamo et al., 2017; Kawana et al., 2017; Nag et al., 2017). We conclude that it is a reasonable hypothesis that myosin missense HCM mutations weaken the SRX-IHM state, resulting in more functionally accessible myosin heads for actin interaction, thus causing the hyper-contractility observed for HCM clinically.

In support of this model, several positively-charged mesa HCM residues, R249Q, H251N, and R453C, significantly weaken the binding of proximal S2 to sS1 in biochemical experiments with purified myosin fragments, and D906G on the proximal S2 that is near the interface with the mesa in the human ß-cardiac myosin folded-back homology models has the same effect (Adhikari et al., 2016; Nag et al., 2017). Interestingly, R403Q has no effect on the affinity of S2 for sS1 (Nag et al., 2017), while here we show that it reduces the amount of SRX in muscle fibers.

So what is R403Q doing at the molecular level? R403Q is in a unique position on the myosin surface, residing at the tip of a pyramid formed by three interacting faces, the actin binding face, the mesa, and the PHHIS (Nag et al., 2017; Trivedi et al., 2017). R403Q does not seem to have a significant effect on actin interaction (Nag et al., 2015), but it could be weakening the complex by weakening the blocked head PHHIS interaction with the free head converter(Nag et al., 2017; Trivedi et al., 2017). Arg403 is also in a position to possibly bind to part of MyBP-C(Nag et al., 2017; Trivedi et al., 2017) and the R403Q mutation may weaken a MyBP-C-SRX complex, and in this way shift the equilibrium away from the SRX in fibers.

Similarly, it is not known which of these various interactions the mesa HCM mutation R663H is affecting, but we show here that it reduces the level of SRX in fibers. Biochemical experiments involving the effects of R403Q and R663H on the interactions between these human ß-cardiac sarcomeric protein players will be necessary to elucidate the molecular mechanism of how these mutations reduce the SRX in the fiber.

The small molecule mavacamten, a cardiac inhibitor in phase 2 clinical trials, decreases contractility and suppresses the development of hypertrophy and fibrosis in HCM mouse models (Green et al., 2016). How is this molecule affecting the detailed structure of the myosin active site? The presence of ATP or ADP.Pi analogs in the active site of myosin causes the switch elements to adopt a closed state, which in-turn has been inferred to stabilize the helical order of the thick filament(Xu et al., 1999; Xu et al., 2003; Zoghbi et al., 2004). Blebbistatin, a small molecule ATPase inhibitor, stabilizes a closed switch-2 state and has also been shown by electron microscopy (Zhao et al., 2008) and fluorescence-based studies (Kampourakis et al., 2018)to stabilize the IHM state of myosin. Conversely, omecamtivmecarbil (OM), a cardiac activator in phase 3 clinical trials, has been shown to stabilize the open state of myosin on the cardiac thick filament (Kampourakis et al., 2018). Along similar lines, mavacamten has been previously shown to reduce the basal release rates of ADP and Pi (Kawas et al., 2017), possibly by holding the switch elements in a closed state, which ultimately stabilizes the myosin heads in the SRX-IHM state. The high-resolution detailed structural basis of formation of the SRX-IHM state is an intriguing, open question in the field.It is crucial now to obtain a high-resolution crystallographic or cryo-EM structure of the SRX-IHM.

We provide here the first evidence at the biochemical level that the cardiac SRX can occur with nothing more than an HMM-like molecule containing two S1 heads and the proximal part of the coiled-coil tail, and that the level of SRX decreases with increasing ionic strength, consistent with many charge-charge interactions stabilizing the IHM. Our work demonstrates that the tail region of myosin is necessary to fully stabilize its SRX conformation. Our 2-hep HMM, however, shows SRX behavior without the proximal tail, which may indicate that the PHHIS-converter association (Kawana et al., 2017; Nag et al., 2017; Trivedi et al., 2017), as well as other important interactions (Alamo et al., 2017) may be enough to hold the heads in a less stable SRX. But even sS1 shows a 15-20% population of SRX, which increases to ∼50% SRX in the presence of mavacamten. We think it likely, therefore, that a small part of the population of independent sS1 molecules in solution exist in a conformation that is similar to that stabilized in the IHM structure (Fig. S4), and the binding of mavacamten shifts the equilibrium of sS1 heads in solution toward the structure of the heads in the SRX-IHM state. Structural studies will be necessary to assess this possibility further We hypothesize that the sS1 SRX conformation will have its light-chain binding helix tilted a bit further in the pre-stroke direction than normally achieved by the relaxed DRX pre-stroke state(Fig. S4). Although a different conformation of the myosin head, this direction of change of lever arm orientation is the same as expected for that of the actin-bound force-producing myosin head under load in contracting muscle, where the light-chain bound lever arm is pulled backward by the load and nucleotide release is slowed(Liu et al., 2018).

The SRX off-state to free head on-state transition of the myosin heads in muscle is thought to be modulated by various factors One possible factor is the stress sensed by the thick filament as a function of the number of heads that are in the on-state and interacting with actin. This stress-sensing mechanism may be important to maintain the helical order of the myosin heads on the thick filament, and increases in the stress level leads to a destabilization of the SRX-IHM and subsequent opening of the heads(Ait-Mou et al., 2016; Irving, 2017; Linari et al., 2015; Reconditi et al., 2017). Thus, there is cross-talk between the global thick filament structure and the conformation of the myosin heads in the SRX-IHM. It has been suggested that this tension sensing is likely to explain the Frank-Starling effect (Ait-Mou et al., 2016; Reconditi et al., 2017), long debated with regard to its mechanism of action(Solaro, 2007).

Other factors modulating the SRX off-state to free head on-state transition could be phosphorylation of the regulatory light chain (Kampourakis and Irving, 2015; Kampourakis et al., 2016; Nag et al., 2017; Trivedi et al., 2017), and/or phosphorylation of MyBP-C (Kampourakis et al., 2014; Nag et al., 2017; Trivedi et al., 2017). There are reported phosphorylation sites on the cardiac ELC as well (Huang and Szczesna-Cordary, 2015), whose possible role in the stabilization of the SRX is unknown. A great deal of work needs to be done to determine what factors modulate the SRX off-state to free head on-state transition.

Our work presents the first structural and biochemical evidence equating IHM to the SRX of myosin and shows that human ß-cardiac myosin HCM mutations in fibers both from a heterozygous piglet and a human heterozygous patient cause a reduction in the SRX. This work opens up avenues for the development of novel therapeutics that specifically target the open-closed equilibrium of the myosin heads. It is conceivable that future cardiac activators can be designed such that they stabilize the open state, while inhibitors like mavacamten stabilize the closed state of the myosin molecule in the thick filament. This forms an innovative basis of modulating cardiac contractility at the molecular level.

## Acknowledgements

The authors thank Drs. Robert McDowell, Leslie Leinwand, Christine Seidman and Hector Rodriguez for useful discussions throughout this work. We thank Dr. Sharlene Day of University of Michigan for providing human R663H and WT tissue samples and the MyoKardia medicinal chemistry group for preparation of mavacamten with a photoactivatable crosslinker. We thank Dr. Roger Craig and Dr. Kyoung-Hwan Lee of University of Massachusetts, Worcester for the invaluable training and guidance during initial phases of the electron microscopy work. We also thank John Perrino of the Stanford Microscopy Facility for technical help with the electron microscopy work, and all members of the Spudich lab for discussions. This work was funded by NIH grants GM33289 and HL117138 (J.A.S.), and AR062279(R.C.). D.V.T is supported by a Stanford Lucile Packard CHRI Postdoctoral Award (UL1 TR001085) and an American Heart Association Postdoctoral Fellowship (17POST33411070). The electron microscopy project described was supported, in part, by ARRA Award Number 1S10RR026780-01 from the National Center for Research Resources (NCRR). Its contents are solely the responsibility of the authors and do not necessarily represent the official views of the NCRR or the National Institutes of Health.This research used the Nikon Imaging center, UCSF; and, resources of the Advanced Photon Source, a U.S. Department of Energy (DOE) Office of Science User Facility operated for the DOE Office of Science by Argonne National Laboratory under Contract No. DE-AC02-06CH11357. This project was supported by grant 9 P41 GM103622 from the National Institute of General Medical Sciences of the National Institutes of Health. The content is solely the responsibility of the authors and does not necessarily reflect the official views of the National Institute of General Medical Sciences or the National Institutes of Health (T.C.I.).

## Author contributions

E.M.G. and J.A.S. conceived and supervised the research. D.V.T., S.S.S.and M.M. performed biochemical experiments with purified human ß-cardiac myosins and analyzed the data. D.V.T., S.S.S. and K.M.R. expressed and purified the proteins. R.L.A., M.H., W.M., H.G and F.L.W., T.C.I and R.C. performed biochemical and structural experiments on porcine and human skinned fibers and analyzed the data. Figures were prepared by R.L.A., D.V.T., S.S.S., E.M.G. and J.A.S. All authors discussed the data as it evolved and reviewed and edited the paper.

## Declaration of Interests

J.A.S. is a founder of Cytokinetics and MyoKardia, a biotechnology company developing small molecules that target the sarcomere for the treatment of inherited cardiomyopathies, and a member of their advisory boards. R.L.A., M.H., and F.L.W. are employees of and own shares in MyoKardia. E.M.G. owns shares in MyoKardia. K.M.R. and R.C. are members of the MyoKardia scientific advisory board.The content is solely the responsibility of the authors and does not necessarily represent the official view of the National Institutes of Health.

## References

Adhikari, A.S., Kooiker, K.B., Sarkar, S.S., Liu, C., Bernstein, D., Spudich, J.A., and Ruppel, K.M. (2016). Early-Onset Hypertrophic Cardiomyopathy Mutations Significantly Increase the Velocity, Force, and Actin-Activated ATPase Activity of Human beta-Cardiac Myosin. Cell Rep 17, 2857–2864.

Ait-Mou, Y., Hsu, K., Farman, G.P., Kumar, M., Greaser, M.L., Irving, T.C., and de Tombe, P.P. (2016). Titin strain contributes to the Frank-Starling law of the heart by structural rearrangements of both thin-and thick-filament proteins. Proc Natl Acad Sci U S A 113, 2306–2311.

Al-Khayat, H.A., Kensler, R.W., Squire, J.M., Marston, S.B., and Morris, E.P. (2013). Atomic model of the human cardiac muscle myosin filament. Proc Natl Acad Sci U S A110, 318–323.

Alamo, L., Qi, D., Wriggers, W., Antonio, Zhu J., Bilbao, A., Gillilan, R.E., Hu, S., and Padron, R. (2016). Conserved intramolecular interactions maintain myosin interacting-heads motifs explaining tarantula muscle super-relaxed state structural basis. J Mol Biol.

Alamo, L., Ware, J.S., Pinto, A., Gillilan, R.E., Seidman, J.G., Seidman, C.E., and Padron, R. (2017). Effects of myosin variants on interacting-heads motif explain distinct hypertrophic and dilated cardiomyopathy phenotypes. Elife6.

Alamo, L., Wriggers, W., Pinto, A., Bartoli, F., Salazar, L., Zhao, F.Q., Craig, R., and Padron, R. (2008). Three-dimensional reconstruction of tarantula myosin filaments suggests how phosphorylation may regulate myosin activity. J Mol Biol384, 780–797.

Blankenfeldt, W., Thoma, N.H., Wray, J.S., Gautel, M., and Schlichting, I. (2006). Crystal structures of human cardiac beta-myosin II S2-Delta provide insight into the functional role of the S2 subfragment. Proc Natl Acad Sci U S A103, 17713–17717.

Burgess, S.A., Yu, S., Walker, M.L., Hawkins, R.J., Chalovich, J.M., and Knight, P.J. (2007). Structures of smooth muscle myosin and heavy meromyosin in the folded, shutdown state. J Mol Biol372, 1165–1178.

Cooke, R. (2011). The role of the myosin ATPase activity in adaptive thermogenesis by skeletal muscle. Biophys Rev3, 33–45.

Geisterfer-Lowrance, A.A., Christe, M., Conner, D.A., Ingwall, J.S., Schoen, F.J., Seidman, C.E., and Seidman, J.G. (1996). A mouse model of familial hypertrophic cardiomyopathy. Science272, 731–734.

Geisterfer-Lowrance, A.A., Kass, S., Tanigawa, G., Vosberg, H.P., McKenna, W., Seidman, C.E., and Seidman, J.G. (1990). A molecular basis for familial hypertrophic cardiomyopathy: a beta cardiac myosin heavy chain gene missense mutation. Cell62, 999–1006.

Green, E.M., Wakimoto, H., Anderson, R.L., Evanchik, M.J., Gorham, J.M., Harrison, B.C., Henze, M., Kawas, R., Oslob, J.D., Rodriguez, H.M., et al. (2016). A small-molecule inhibitor of sarcomere contractility suppresses hypertrophic cardiomyopathy in mice. Science 351, 617–621.

Homburger, J.R., Green, E.M., Caleshu, C., Sunitha, M.S., Taylor, R.E., Ruppel, K.M., Metpally, R.P., Colan, S.D., Michels, M., Day, S.M., et al. (2016). Multidimensional structure-function relationships in human beta-cardiac myosin from population-scale genetic variation. Proc Natl Acad Sci U S A 113, 6701–6706.

Hooijman, P., Stewart, M.A., and Cooke, R. (2011). A new state of cardiac myosin with very slow ATP turnover: a potential cardioprotective mechanism in the heart. Biophys J100, 1969–1976.

Huang, W., and Szczesna-Cordary, D. (2015). Molecular mechanisms of cardiomyopathy phenotypes associated with myosin light chain mutations. J Muscle Res Cell Motil36, 433–445.

Irving, M. (2017). Regulation of Contraction by the Thick Filaments in Skeletal Muscle. Biophys J113, 2579–2594.

Jung, H.S., Billington, N., Thirumurugan, K., Salzameda, B., Cremo, C.R., Chalovich, J.M., Chantler, P.D., and Knight, P.J. (2011). Role of the tail in the regulated state of myosin 2. J Mol Biol408, 863–878.

Jung, H.S., Komatsu, S., Ikebe, M., and Craig, R. (2008). Head-head and head-tail interaction: a general mechanism for switching off myosin II activity in cells. Mol Biol Cell19, 3234–3242.

Kampourakis, T., and Irving, M. (2015). Phosphorylation of myosin regulatory light chain controls myosin head conformation in cardiac muscle. J Mol Cell Cardiol85, 199–206.

Kampourakis, T., Sun, Y.B., and Irving, M. (2016). Myosin light chain phosphorylation enhances contraction of heart muscle via structural changes in both thick and thin filaments. Proc Natl Acad Sci U S A113, E3039–3047.

Kampourakis, T., Yan, Z., Gautel, M., Sun, Y.B., and M., I. (2014). Myosin binding protein-C activates thin filaments and inhibits thick filaments in heart muscle cells. Proc Natl Acad Sci U S A111, 18763–18768.

Kampourakis, T., Zhang, X., Sun, Y.B., and Irving, M. (2018). Omecamtiv mercabil and blebbistatin modulate cardiac contractility by perturbing the regulatory state of the myosin filament. J Physiol596, 31–46.

Kawana, M., Sarkar, S.S., Sutton, S., Ruppel, K.M., and Spudich, J.A. (2017). Biophysical properties of human beta-cardiac myosin with converter mutations that cause hypertrophic cardiomyopathy. Sci Adv3, e1601959.

Kawas, R.F., Anderson, R.L., Ingle, S.R.B., Song, Y., Sran, A.S., and Rodriguez, H.M. (2017). A small-molecule modulator of cardiac myosin acts on multiple stages of the myosin chemomechanical cycle. J Biol Chem292, 16571–16577.

Linari, M., Brunello, E., Reconditi, M., Fusi, L., Caremani, M., Narayanan, T., Piazzesi, G., Lombardi, V., and Irving, M. (2015). Force generation by skeletal muscle is controlled by mechanosensing in myosin filaments. Nature528, 276–279.

Liu, C., Kawana, M., Song, D., Ruppel, K.M., and Spudich, J. (2018). Controlling load-dependent contractility of the heart at the single molecule level. bioRxiv.

Maron, M., Ashley, E., Blok, T., Evanchik, M., Lambing, J., Lester, S., Mahaffey, K., Markova, S., Owens, A., Ritter, J., et al. (2016). Abstract 16842: Obstructive Hypertrophic Cardiomyopathy: Initial Single Ascending Dose Data in Healthy Volunteers and Patients. Circulation 134, A16842–A16842.

McNamara, J.W., Li, A., Dos Remedios, C.G., and Cooke, R. (2015). The role of super-relaxed myosin in skeletal and cardiac muscle. Biophys Rev7, 5–14.

McNamara, J.W., Li, A., Smith, N.J., Lal, S., Graham, R.M., Kooiker, K.B., van Dijk, S.J., Remedios, C.G., Harris, S.P., and Cooke, R. (2016). Ablation of cardiac myosin binding protein-C disrupts the super-relaxed state of myosin in murine cardiomyocytes. J Mol Cell Cardiol94, 65–71.

Moore, J.R., Leinwand, L., and Warshaw, D.M. (2012). Understanding Cardiomyopathy Phenotypes Based on the Functional Impact of Mutations in the Myosin Motor. Circulation Research111, 375–385.

Naber, N., Cooke, R., and Pate, E. (2011). Slow myosin ATP turnover in the super-relaxed state in tarantula muscle. J Mol Biol411, 943–950.

Nag, S., Sommese, R.F., Ujfalusi, Z., Combs, A., Langer, S., Sutton, S., Leinwand, L.A., Geeves, M.A., Ruppel, K.M., and Spudich, J.A. (2015). Contractility parameters of human beta-cardiac myosin with the hypertrophic cardiomyopathy mutation R403Q show loss of motor function. Sci Adv1, e1500511.

Nag, S., Trivedi, D.V., Sarkar, S.S., Adhikari, A.S., Sunitha, M.S., Sutton, S., Ruppel, K.M., and Spudich, J.A. (2017). The myosin mesa and the basis of hypercontractility caused by hypertrophic cardiomyopathy mutations. Nat Struct Mol Biol.

Nag, S., Trivedi, D.V., Sarkar, S.S., Sutton, M.S., Ruppel, K.M., and Spudich, J.A. (2016). Beyond the myosin mesa: a potential unifying hypothesis on the underlying molecular basis of hyper-contractility caused by a majority of hypertrophic cardiomyopathy mutations. bioRxiv.

Nogara, L., Naber, N., Pate, E., Canton, M., Reggiani, C., and Cooke, R. (2016). Spectroscopic Studies of the Super Relaxed State of Skeletal Muscle. PLoS One11, e0160100.

Previs, M.J., Prosser, B.L., Mun, J.Y., Previs, S.B., Gulick, J., Lee, K., Robbins, J., Craig, R., Lederer, W.J., and Warshaw, D.M. (2015). Myosin-binding protein C corrects an intrinsic inhomogeneity in cardiac excitation-contraction coupling. Sci Adv

Reconditi, M., Caremani, M., Pinzauti, F., Powers, J.D., Narayanan, T., Stienen, G.J., Linari, M., Lombardi, V., and Piazzesi, G. (2017). Myosin filament activation in the heart is tuned to the mechanical task. Proc Natl Acad Sci U S A114, 3240–3245.

Seidman, C.E., and Seidman, J.G. (2000). Hypertrophic cardiomyopathy. In The Metabolic and Molecular Bases of Inherited Disease, C.R. Scriver, A.L. Beaudet, D. Valle, W.S. Sly, K.W. Childs, and B. Vogelstein, eds. (McGraw-Hill), pp. 5532–5452.

Seidman, J.G., and Seidman, C.E. (2001). The Genetic Basis for Cardiomyopathy: from Mutation Identification to Mechanistic Paradigms. Cell 104, 557–567.

Solaro, R.J. (2007). Mechanisms of the Frank-Starling law of the heart: the beat goes on. Biophys J93, 4095–4096.

Spudich, J.A. (2014). Hypertrophic and dilated cardiomyopathy: four decades of basic research on muscle lead to potential therapeutic approaches to these devastating genetic diseases. Biophys J106, 1236–1249.

Spudich, J.A. (2015). The myosin mesa and a possible unifying hypothesis for the molecular basis of human hypertrophic cardiomyopathy. Biochem Soc Trans43, 64–72.

Spudich, J.A., Aksel, T., Bartholomew, S.R., Nag, S., Kawana, M., Yu, E.C., Sarkar, S.S., Sung, J., Sommese, R.F., Sutton, S., et al. (2016). Effects of hypertrophic and dilated cardiomyopathy mutations on power output by human beta-cardiac myosin. J Exp Biol 219, 161–167.

Stewart, M.A., Franks-Skiba, K., Chen, S., and Cooke, R. (2010). Myosin ATP turnover rate is a mechanism involved in thermogenesis in resting skeletal muscle fibers. Proc Natl Acad Sci U S A107, 430–435.

Toyoshima, Y.Y., Kron, S.J., McNally, E.M., Niebling, K.R., Toyoshima, C., and Spudich, J.A. (1987). Myosin subfragment-1 is sufficient to move actin filaments in vitro. Nature 328, 536–539.

Trivedi, D.V., Adhikari, A.S., Sarkar, S.S., Ruppel, K.M., and Spudich, J.A. (2017). Hypertrophic cardiomyopathy and the myosin mesa: viewing an old disease in a new light. Biophys Rev.

Tyska, M.J., Hayes, E., Giewat, M., Seidman, C.E., Seidman, J.G., and Warshaw, D.M. (2000). Single-molecule mechanics of R403Q cardiac myosin isolated from the mouse model of familial hypertrophic cardiomyopathy. Circ Res86, 737–744.

Wendt, T., Taylor, D., Messier, T., Trybus, K.M., and Taylor, K.A. (1999). Visualization of head-head interactions in the inhibited state of smooth muscle myosin. J Cell Biol147, 1385–1390.

Wendt, T., Taylor, D., Trybus, K.M., and Taylor, K. (2001). Three-dimensional image reconstruction of dephosphorylated smooth muscle heavy meromyosin reveals asymmetry in the interaction between myosin heads and placement of subfragment 2. Proc Natl Acad Sci U S A98, 4361–4366.

Wilson, C., Naber, N., Pate, E., and Cooke, R. (2014). The myosin inhibitor blebbistatin stabilizes the super-relaxed state in skeletal muscle. Biophys J107, 1637–1646.

Wilson, W.S., Criley, J.M., and Ross, R.S. (1967). Dynamics of left ventricular emptying in hypertrophic subaortic stenosis. A cineangiographic and hemodynamic study. Am Heart J 73, 4–16.

Woodhead, J.L., Zhao, F.Q., Craig, R., Egelman, E.H., Alamo, L., and Padron, R. (2005). Atomic model of a myosin filament in the relaxed state. Nature436, 1195–1199.

Xu, S., Gu, J., Rhodes, T., Belknap, B., Rosenbaum, G., Offer, G., White, H., and Yu, L.C. (1999). The M.ADP.Pi state is required for helical order in the thick filaments of skeletal muscle. Biophys J77, 2665–2676.

Xu, S., Offer, G., Gu, J., White, H.D., and Yu, L.C. (2003). Temperature and ligand dependence of conformation and helical order in myosin filaments. Biochemistry42, 390–401.

Zhao, F.Q., Padron, R., and Craig, R. (2008). Blebbistatin stabilizes the helical order of myosin filaments by promoting the switch 2 closed state. Biophys J95, 3322–3329.

Zoghbi, M.E., Woodhead, J.L., Craig, R., and Padron, R. (2004). Helical order in tarantula thick filaments requires the “closed” conformation of the myosin head. J Mol Biol 342, 1223–1236.

Zoghbi, M.E., Woodhead, J.L., Moss, R.L., and Craig, R. (2008). Three-dimensional structure of vertebrate cardiac muscle myosin filaments. Proc Natl Acad Sci U S A 105, 2386–2390.

